# The connectome predicts resting state functional connectivity across the Drosophila brain

**DOI:** 10.1101/2020.12.11.422105

**Authors:** Maxwell H. Turner, Kevin Mann, Thomas R. Clandinin

**Author notes:** Equal contribution (Maxwell H. Turner, Kevin Mann).

## Abstract

Connectomic datasets have emerged as invaluable tools for understanding neural circuits in many systems. What constraints does the connectome place on information processing and routing in a large scale neural circuit? For mesoscale brain networks, the relationship between cell and synaptic level connectivity and brain function is not well understood. Here, we use data from the Drosophila connectome in conjunction with whole-brain *in vivo* imaging to relate structural and functional connectivity in the central brain. We find that functional connectivity is strongly associated with the strength of both direct and indirect anatomical pathways. We also show that some brain regions, including the mushroom body and central complex, show considerably higher functional connectivity to other brain regions than is predicted based on their direct anatomical connections. We find several key topological similarities between mesoscale brain networks in flies and mammals, revealing conserved principles relating brain structure and function.

## Introduction

Anatomical connectivity can constrain both a neural circuit’s function and its underlying computation. This principle has been demonstrated for many small, defined neural circuits. For example, connectome reconstructions have informed models for direction selectivity in the vertebrate retina (Helmstaedter et al. 2013; Kim et al. 2014) as well as the Drosophila visual system (Takemura et al. 2017a). In these cases, the circuit in question is relatively compact, well-defined, and has known functions. However, whether and how the connectome constrains global properties of large-scale networks, across multiple brain regions or the entire brain, is unknown. As the availability of partial or complete connectomes expands to more systems and species (Scheffer et al. 2020; Cook et al. 2019; Kasthuri et al. 2015; Markram et al. 2015; Abbott et al. 2020) it becomes critical to understand how this detailed anatomical information can inform our understanding of large scale circuit function and computation (Bargmann and Marder 2013; Meinertzhagen 2018).

An illuminating approach to understanding large-scale structure-function relationships in the brain has emerged in human neuroscience, where macro-scale structural connectivity patterns can be related to functional connectivity using noninvasive imaging or electrical recording techniques (for reviews, see Bullmore and Sporns 2009; Suárez et al. 2020). Here, structural connectivity typically refers to the density of axon fibers connecting two (cortical) brain regions, and functional connectivity refers to the patterns of inter-region correlations in measures of brain activity under task-based or resting state conditions. Several of these studies have shown that functional connectivity can be predicted by structural connectivity to some degree (Honey et al. 2009; Honey et al. 2007; Hermundstad et al. 2013; Suárez et al. 2020). However, the apparent structure-function correlation is limited, which is partially the result of not knowing the structural connectivity of the human brain at single cell resolution. How well can functional connectivity be predicted given a complete anatomical reconstruction?

The Drosophila nervous system is a powerful model for understanding circuit function at the level of identified cell types and synapses. In this system, the wiring diagram is highly stereotyped from animal to animal and many studies have characterized neural activity and behavior in a variety of contexts. As a result, there is a deep understanding of many individual circuits and brain regions. However, how circuit activity is coordinated across brain regions remains incompletely understood. We have previously developed techniques for measuring neural activity across the entire Drosophila central brain while assigning functional responses to the brain regions defined in a common brain atlas (Mann et al. 2017). These data, combined with the completion of the Drosophila central brain connectome (Scheffer et al. 2020) allow us to explore the relationship between synapse-level structural connectivity and brain-wide functional connectivity.

Here, we ask several related questions about the relationship between structural connectivity and functional connectivity in the Drosophila central brain: (1) How is resting state functional connectivity shaped by structural connectivity in the central brain? (2) To what extent does detailed, synapse-level connectivity determine mesoscale functional interactions in the brain? And (3) What role do higher order network interactions play in determining functional connectivity? We find a strong relationship between resting state functional connectivity and direct region-to-region structural connectivity. Surprisingly, including information about the number of synaptic connections does not improve the correlation between structural and functional connectivity beyond a connectivity metric that only considers cell number. We also provide evidence that indirect pathways in the brain differentially shape functional connectivity, with a small subset of regions being strongly dependent on indirect connections. Throughout this work we observe features of structural and functional networks in Drosophila that are strikingly similar to those seen previously in mammalian cortex, including in the human brain. Given the vast anatomical and functional differences between Drosophila and mammalian nervous systems, these observations suggest general principles that govern brain structure, function and the relationship between the two.

## Results

In order to study global properties of brain networks, we focused on connectivity among regions (“mesoscale connectivity”), a scale at which we could both reliably register neural activity signals across individuals and make comparisons to network organization in other systems. To characterize mesoscale connectivity, we established methods for comparing structural and functional connectivity in a common anatomical space. We used data from the Drosophila “hemibrain” connectome (Scheffer et al. 2020). This dataset contains most of the right half of the central brain of a single fly, and includes ~25,000 traced cells and ~10 million presynaptic active zones. This connectome has been reconstructed with high accuracy, with over 90% of presynaptic sites (T-Bars) assigned to traced neurons (for details of the hemibrain and its reconstruction, see Scheffer et al. 2020). To measure mesoscale functional connectivity, we used pan-neuronal, *in vivo* calcium imaging of the central brain (Mann et al. 2017) in the absence of sensory stimuli. Each brain (n=20) comprised ~500,000 voxels, and was imaged at an isotropic resolution of 3 *μ*m. These voxels were then assigned to anatomical regions. From these measurements, we computed characteristic correlations among brain regions that we refer to as the resting state functional connectivity. Finally, we compare both structural and functional datasets in a common anatomical space (Ito et al. 2014). We included the central brain regions with at least half of their volume included in the hemibrain. In total this analysis includes 36 brain regions (see Supplementary table 1 for list of regions) that follow the naming convention outlined in Ito et al. 2014.

### Characterizing structural connectivity

The mesoscale connectivity network is defined by the connection strength between every pair of regions. We used two primary metrics of connection strength (Fig. 1A,B). First, the cell count connectivity from region A to region B was defined as the total number of neurons with at least one input (post-) synapse in region A and at least one output (pre-) synapse in region B. Second, the T-Bar connectivity was defined as the total number of presynaptic T-Bars in region B that belong to a neuron with at least one input in region A.

**Figure 1:**
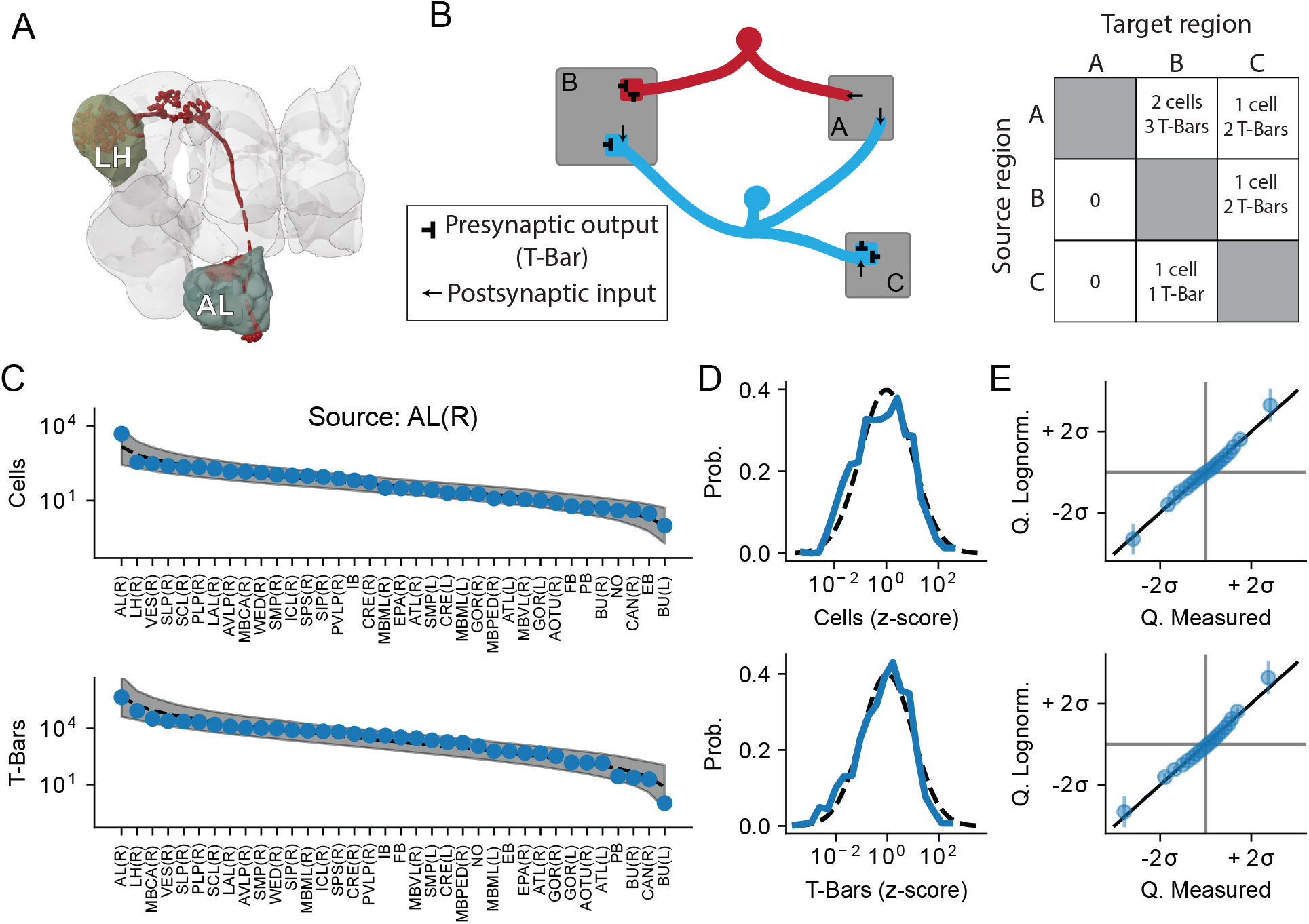
Characterizing mesoscale connectivity using the hemibrain connectome. (A) Top: Portion of the central brain included in the hemibrain connectome used for this analysis, with highlighted example regions. Shown in red is an example of a neuron which projects from the antenal lobe (AL) to the lateral horn (LH) (B) Schematic showing how inter-region connectivity is quantified. For example, the connection from region A to region B has 2 cells and 3 T-Bars. (C) For an example region (the right antennal lobe - AL(R)) the number of outgoing connections to all regions included in this analysis, ordered from highest to lowest connection strength. Top: inter-region connectivity based on cell count, bottom: inter-region connectivity based on presynaptic T-Bars. Data are in blue, dashed line and shaded region indicate the mean ±2*σ* of a log-normal distribution with mean and standard deviation measured from the data. (D) Inter-region connectivity strength follows a log-normal distribution. Top: Outgoing cell count connectivity from each region was z-scored within a region and combined to produce a distribution of connection strengths across all regions (solid blue line). Dashed black line shows, for reference, a log-normal distribution with mean and standard deviation matched to the data. Bottom: same as top, but using T-Bar count as the connectivity metric. (E) Top: Q-Q plot to compare the distribution of observed cell count connectivity (horizontal axis) to a best-fitting log-normal distribution (vertical axis). Bottom: same as top, but using T-Bar count as the connectivity metric.

The outgoing connectivity distributions, for each metric, are shown for an example region - the right antennal lobe (AL(R), Fig. 1C). The outgoing connectivity strengths span several orders of magnitude for both the cell count and T-Bar metrics. For this example source region, the strongest downstream partner, the right lateral horn (LH(R)), has several thousand cell connections, while the weakest, the left bulb (BU(L)) receives only a few. Interestingly, the outgoing connections closely follow a log-normal distribution of connectivity strengths. A similar pattern was observed for every brain region for both the cell count and T-Bar connectivity metrics (p>0.05, KS test against a log-normal distribution for each region, see Fig. 1D,E & Fig. S1). A similar log-normal distribution of connectivity strengths has been shown for inter-region connectivity in mammalian cortex, including in mouse (Wang et al. 2012; Oh et al. 2014) and monkey (Markov et al. 2014).

### The connectome predicts resting state functional connectivity

We used our whole brain functional imaging data to measure resting state correlations among brain regions. We registered our brain volumes to a common anatomical atlas (Fig. 2A) and extracted the average fluorescence signal for each brain region over a period of approximately 25 minutes (Fig. 2B). We then computed the correlation between these signals for each pair of regions (Fig. 2C), a measure of their functional connectivity, and averaged these correlations across animals (Fig. 2D). Note that the functional correlation values here have been Fisher z-transformed to facilitate estimating statistics across animals. The structural connectivity matrix is shown in Fig. 2E, using the cell count connectivity metric (the T-Bar connectivity matrix is very similar, r=0.90).

**Figure 2:**
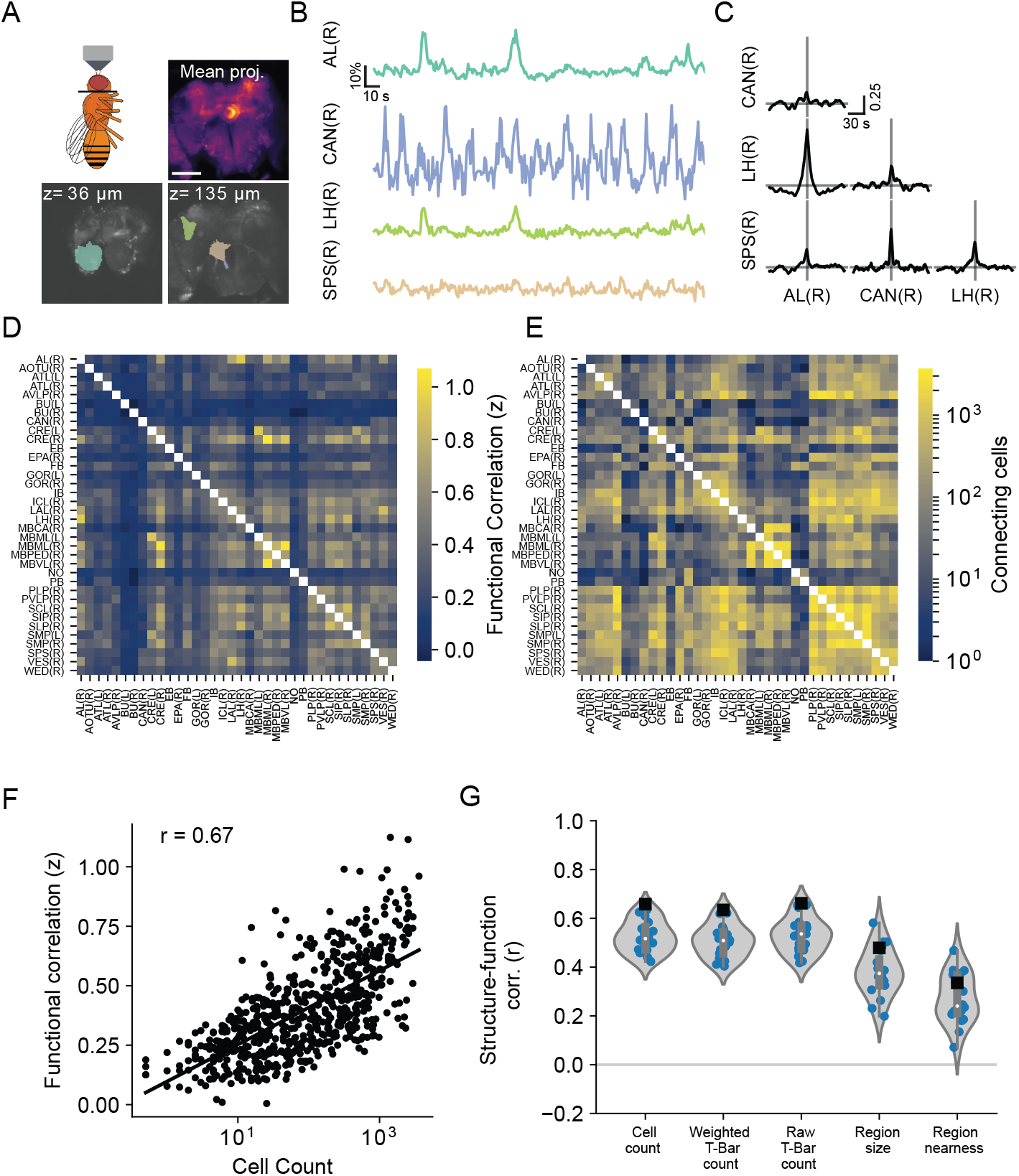
Structural connectivity revealed by the connectome predicts resting state functional connectivity in the central brain. (A) Overview of functional imaging acquisition. Top left: the central brain is imaged from the anterior of the fly head. Top right: mean projection showing central brain expressing GCaMP6s. Scale bar is 100 *μm*. Bottom: single planes through the central brain showing brain regions from an anatomical atlas that has been aligned to *in vivo* data. (B) Example traces showing GCaMP6s responses of representative brain regions. (C) Table of cross-correlograms for the pairwise combinations of the four highlighted brain regions in (A,B). (D) Average correlation matrix across n=20 flies, each entry in the matrix is the Fisher transformed correlation value for that pair of regions. (E) Heatmap showing cell count connectivity between every pair of regions included in this analysis. Regions are ordered alphabetically. (F) The log-transformed cell count connectivity between a pair of brain regions is strongly correlated with the functional correlation between those regions. (G) For each anatomical connectivity metric, the distribution of structure-function correlation values for individual flies (blue points and violin plot) and the correlation between structure and the across-animal average functional connectivity matrix (black square).

By visual inspection, the functional connectivity in the central brain appears similar to the mesoscale connectivity revealed by the connectome (compare Fig. 2D and 2E). To quantify this relationship, we plotted the functional correlation between each pair of regions against their structural connectivity (Fig. 2F). This analysis reveals a positive correlation between structural and functional connectivity in the central brain (Pearson r=0.66). However, the connectome contains connectivity information that is not captured by the cell count metric. Does including more detailed synaptic connectivity increase the correlation between structural and functional connectivity? To address this, we also examined how other mesoscale connectivity metrics predict functional connectivity (Fig. 2G).

In addition to the “Cell count” and “T-Bar count” connectivity metrics described above, we also examined the “Weighted synapse count” as defined in Scheffer et al. 2020 - i.e. the T-Bar count is weighted by the fraction of postsynapses onto each cell that are within the source region. Surprisingly, the metrics that include information about synapse number are no more predictive of functional connectivity than cell count alone (Fig. 2G). Other anatomical features, like the sizes of each region and the physical distance between regions, are significantly correlated with functional connectivity, but are less predictive than the connectome-derived metrics (Fig. 2G). The correlation between region size or distance and functional connectivity likely reflects the fact that larger regions contain more cells and synapses than smaller regions and that regions that are nearer to one another tend to have stronger anatomical connections (data not shown). However, when we divided the cell count connectivity by the region size, we found that this normalized structural connectivity measure was still able to predict functional connectivity (Fig. S2). Thus, the relationship between connectome-derived connectivity and functional connectivity is not the result of binning the brain into unequally sized regions.

Importantly, we found that the presence of relatively small brain regions did not preclude the accurate estimation of structure-function correlations (Fig. S3). We also found that connectome reconstruction completeness was unrelated to functional connectivity, and that reconstruction incompleteness did not result in lower cell count connectivity metrics (Fig. S4). The results of these analyses suggest that these data contain the requisite accuracy to reliably capture mesoscale structure-function relationships.

### Graph features of structural and functional connectivity networks

To further examine correspondences between the structural and functional networks, we used the connectivity matrices in Fig. 2 to construct graph representations of each network. In these graphs, each node corresponds to a brain region, and each edge corresponds to the connection between two regions (Bassett et al. 2018). For the structural graph, we symmetrized the connectivity matrix by averaging the outgoing and incoming (i.e. A to B and B to A) connections. Each node of the graph and the top 100 strongest edges of the structural and functional networks are shown in Fig. 3A,B. The size of each node corresponds to its degree (defined as the sum of the weights across all incoming and outgoing connections associated with that node). Graph features in the structural and functional networks are highly correlated, including the node degree (Fig. 3C, Pearson r=0.64) and the node clustering coefficient, which measures how connected a node’s neighbors are (Fig. 3D, Pearson r=0.64). However, there are discrepancies between these two graphs, for example, the AVLP(R) has a relatively low degree and clustering coefficient in the functional graph compared to its structural graph. Conversely, regions like the MBML(R) and LH(R) have relatively high degrees and clustering coefficients compared to their respective structural graphs. These results suggest that the ability to predict features of functional networks from direct anatomical connections is limited, and raises the possibility that indirect connections may also play an important role (see Fig. 4).

**Figure 3:**
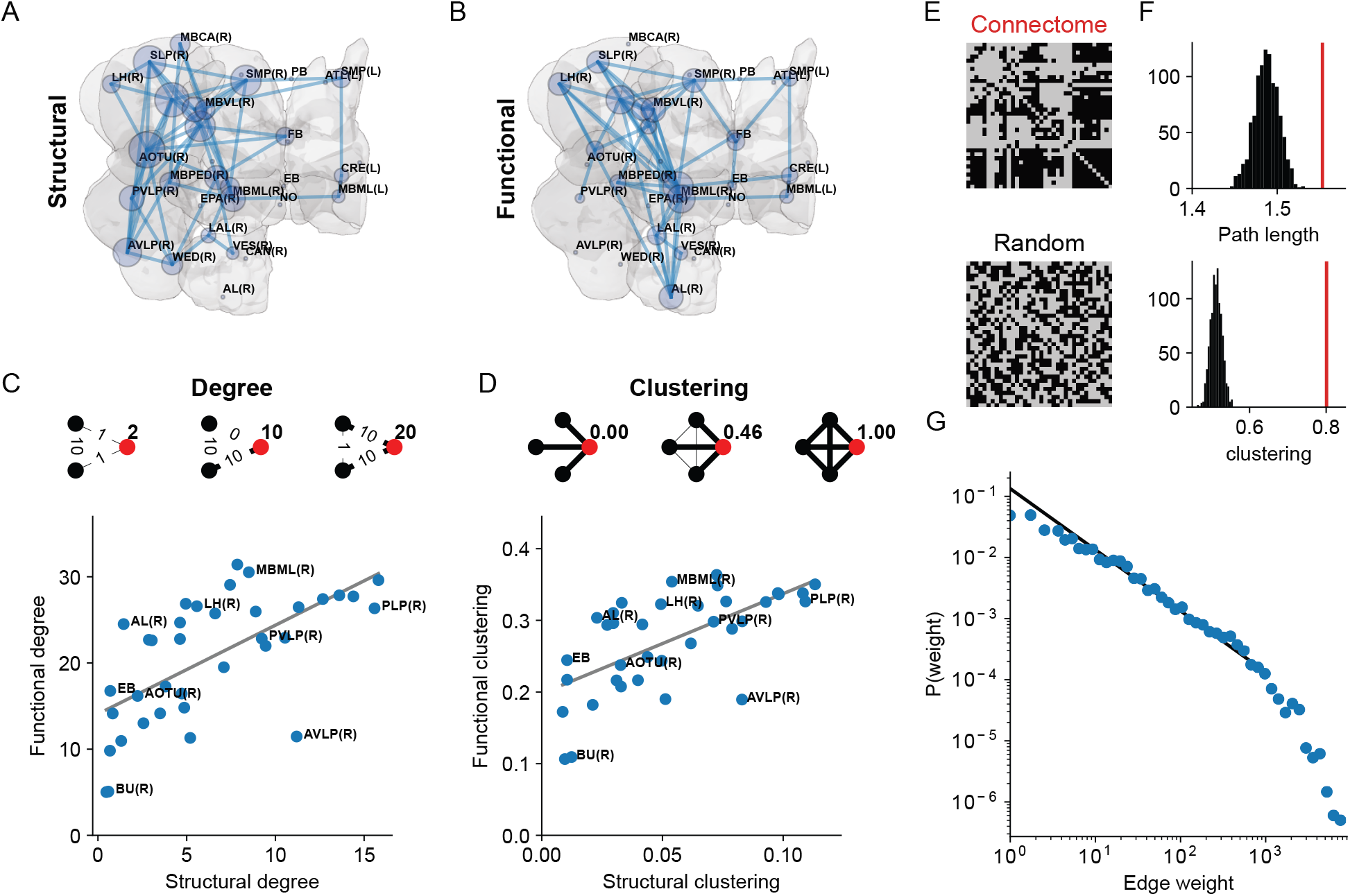
Graph features are shared between structural and functional connectivity networks. (A) graph representation of the connectome-derived structural network. Each node corresponds to a brain region, and the size of each node represents the degree of that node. The top 100 strongest edges are shown as lines connecting nodes. (B) graph representation of the functional network. (C) Top: schematic illustrating the degree of a node. In each graph example, the degree of the red node is represented by the bold number to its right. A high degree indicates a node with many strong connections (both incoming and outgoing). Bottom: The node degree for the structural graph is correlated to the node degree of the functional graph (Pearson r=0.64). (D) Top: schematic illustrating the clustering coefficient of a node. A high clustering coefficient indicates a node whose neighbors are highly connected. Bottom: The clustering coefficients in the structural network are correlated to those of the functional network (Pearson r=0.64). (E) Thresholded structural connectivity matrix and random connectivity matrix with a matched connection probability (F) Top: Distribution of average path lengths across 1000 random graphs (black distribution) and average path length for the measured structural graph (red line). Bottom: same as top but for the average graph clustering coefficient. (G) The distribution of edge weights in the cell-count connectivity network follows a power law distribution for most edge weights. Black line shows power law scaling, p ∝ *w*^−1^, blue points are edge weights from the measured structural connectivity.

**Figure 4:**
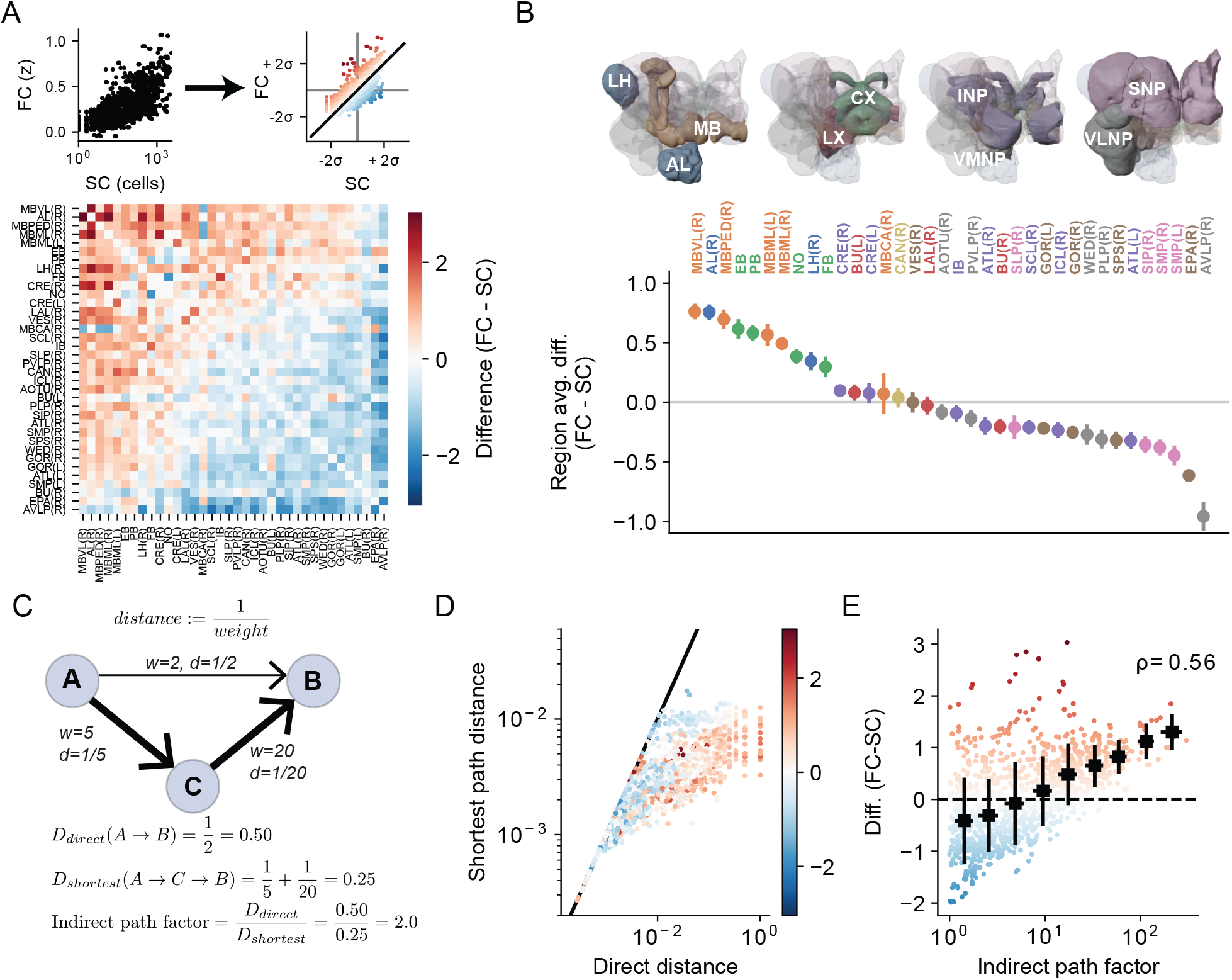
Structure-function relationships in the central brain vary by major brain regions because of indirect connections. (A) Top: We z-scored both structural connectivity (SC) and functional connectivity (FC). Bottom: Difference matrix, showing, for each pair of regions, the difference between FC and SC. Positive (red) values correspond to region pairs that have relatively high FC given their SC, and negative (blue) values correspond to region pairs that have relatively low FC. Regions have been sorted by the region-average difference in descending order. (B) FC-SC differences vary across brain super-regions, with the highest differences clustered in the AL/LH, Central Complex, and Mushroom Body regions. Each point corresponds to the mean ± s.e.m., for that region, across n=20 flies. Points are sorted in descending order of FC-SC difference, and colored according to their membership in super-regions, which are illustrated above. (C) Illustration of direct vs. indirect anatomical pathways that could influence functional connectivity. We define the distance between two nodes as the inverse of that edge’s weight. To quantify the relative strength of indirect pathways, we define an indirect path factor as the ratio of the direct path distance to the shortest path distance. (D) For each pair of regions, the shortest path distance is less than or equal to the direct path distance. Points are color-coded by the FC-SC difference, as in (A). Note that points further from the line of unity tend to have positive FC-SC differences. (E) The strength of indirect pathways in the brain accounts for some of the discrepancy between direct pathway structural connectivity and measured functional connectivity. *ρ* is the Spearman rank order correlation coefficient.

We also investigated whether the structural network shares topological features observed in large scale networks in the human brain. In particular we asked whether the mesoscale network shows properties characteristic of (1) a small-world network and (2) a scale-free network. A small-world network is characterized by strong clustering of nodes and relatively short path lengths through the network, even in the absence of dense connections (Sporns et al. 2004; Sporns and Zwi 2004; Bassett and Bullmore 2006). To determine whether the mesoscale network shows these features, we thresholded the structural connectivity matrix to produce a binary adjacency matrix (Fig. 3E, top). We then generated 1000 random graphs with a connection probability matched to that of the measured adjacency matrix (Fig. 3E, bottom). For both the measured graph and the random graphs, we measured the average path length (Fig. 3F, top) and clustering coefficient (Fig. 3F, bottom). The red vertical line indicates the measured adjacency matrix, while the black distribution shows the computed values for random graphs. As is typical of small-world networks, including connectivity networks in human cortex (Bullmore and Sporns 2009), the Drosophila structural network has a much higher clustering coefficient than equivalent random graphs (56% higher on average), but only slightly longer path length (4% longer on average, see also Shih et al. 2015). A scale-free network is one in which the distribution of node degree or connection weights follows a power law distribution. This structure has been observed in human functional networks (Eguíluz et al. 2005; Bullmore and Bassett 2011). For the vast majority of edges, and over more than 2 orders of magnitude of edge weights, the edge weight distribution of the Drosophila structural network follows a power law decay (Fig. 3G). We note that for very weak edges as well as very strong edges, the data deviate from power law scaling (see Scheffer 2020 for a similar analysis on cell-to-cell connectivity). Taken together, this analysis of the structural connectivity indicates that the Drosophila mesoscale anatomical network shares two salient topological features previously described in human cortical networks.

### Indirect pathways differentially shape structure-function relationships across the brain

Thus far, we have shown that there is a broad correspondence between structural and functional connectivity. However, there are also discrepancies between functional connectivity and what would be predicted based on direct structural connectivity alone (Fig. 3). Are these differences between structural and functional connectivity uniformly distributed throughout the brain, or are they concentrated in certain regions? To answer this question, we z-scored both the structural and functional connectivity and computed the difference (Fig. 4A).

When we averaged the function-structure difference for each region, we found that some brain regions tend to have relatively high functional compared to structural connectivity (Fig. 4B). We color-coded brain regions by the super-region classification scheme in Ito et al. 2014, where a super-region typically contains regions that are nearby to one another and grouped by anatomical boundaries. This analysis revealed heterogeneity in the function-structure difference across brain regions that in some cases corresponded to membership in a super-region. For example, 8 of the 10 regions with the highest average difference (left-most points in Fig. 4B) belong to the mushroom body or central complex. The other two regions are the antennal lobe & lateral horn, which are in distinct super-regions but part of a common circuit (Wilson 2013; Masse et al. 2009; Schultzhaus et al. 2017). On the other hand, regions belonging to the superior neuropils (SNP) and the ventrolateral neuropils (VLNP) had relatively low functional connectivity to other regions given their structural connectivity (right side of Fig. 4B).

What might account for the relatively poor prediction of functional connectivity from structural connectivity in these regions? The analysis above uses *direct* anatomical connections between two regions to define their structural connectivity. But many brain networks include strong indirect pathways between regions (Avena-Koenigsberger et al. 2018). Indeed, this can be a characteristic feature of small-world, scale-free networks (Eguíluz et al. 2005; Sporns et al. 2004; Sporns and Zwi 2004) (Fig. 3). One important consequence of the impact of indirect pathways is that two brain regions may be functionally correlated despite having a weak (or nonexistent) direct connection between them, as multi-synaptic connections could produce correlated neural activity.

To measure the presence of indirect pathways in the inter-region structural network, we first defined the distance between two regions as the inverse of the connectivity weight between them (Fig. 4C). Therefore a pair of regions with a strong connection between them will be separated by a short distance. For each pair of brain regions, A and B, we computed the direct distance (*D*_*direct*_(*A, B*)), which is the distance of the direct pathway between regions A and B. We also computed the shortest path distance (*D*_*shortest*_(*A, B*)) between regions A and B using Dijkstra’s algorithm (Dijkstra 1959). This commonly-used algorithm finds the single shortest path between any two nodes in a (connected) graph. This approach can reveal multi-step, indirect connections between a pair of regions that are stronger than the direct connection between them. Thus, for every pair of regions, the shortest path distance is always less than or equal to the direct distance (Fig. 4D). This analysis shows that connections with higher-than-expected functional connectivity tend to have much longer direct path distances compared to their shortest paths. To quantify this, we defined an indirect path factor, which is the ratio of the direct path distance to the shortest path distance. An indirect path factor of 1.0 indicates a connection where the shortest path is equal in distance to the direct path, and values greater than 1.0 indicate a shortest path which is shorter than the direct path. In this way, the indirect path factor quantifies the relative strength of the primary indirect pathway between a pair of regions. We found that the strength of indirect pathways was positively correlated with the function-structure difference (Spearman’s rank correlation *ρ* = 0.56). This means that regions that showed higher functional connectivity than predicted from direct structural connectivity tend to be connected to other regions by strong indirect pathways.

To further test whether indirect pathways contribute to functional connectivity, we fit linear regression models to predict functional connectivity based on different measures of structural connectivity. We found that a model that included both shortest path distance and direct connectivity outperformed a model that included only direct distance (cell count metric: *r*^2^=0.47 vs. *r*^2^=0.42 for the direct connectivity only model; T-Bars metric: *r*^2^=0.55 vs. *r*^2^=0.42 for the direct connectivity only model) (Fig. S5). Taken together, these results suggest that indirect anatomical pathways in the brain exert an important influence on functional connectivity.

## Discussion

The release of a nearly-complete central brain connectome, combined with the ability to measure whole-brain functional activity in defined central brain regions has allowed us to, for the first time, relate synapse-level structural connectivity to mesoscale functional connectivity across the brain (Mann et al. 2017; Scheffer et al. 2020). We found that cell-level anatomical connectivity provides a strong constraint on resting state functional connectivity (Fig. 2).

While the direct, inter-region structural connectivity was broadly predictive of functional connectivity, we found that indirect pathways could also contribute significantly to functional connectivity (Fig. 4). Indirect connections are especially important for some brain regions in particular, including the mushroom body and central complex, which are associated with multisensory integration and learning (Yagi et al. 2016; Currier and Nagel 2020; Turner-Evans and Jayaraman 2016; Fisher et al. 2019; Kim et al. 2019), suggesting that dense indirect connectedness might be important for these computations. We used one metric of indirect connections between two regions, namely the shortest path distance, but one could imagine other indirect paths also contributing to functional connectivity in a meaningful way. Indeed, recent work developed a graph embedding procedure to predict functional connectivity from structural connectivity in the human brain (Rosenthal et al. 2018; Grover and Leskovec 2016), which allows for the influence of higher order interactions in general. We suspect that, rather than the specific shortest path being of particular importance in shaping functional connectivity between a pair of regions, the shortest path distance is a proxy for more general indirect path connectedness. Disentangling which higher order interactions most strongly shape functional connectivity will be an important task in future work.

The structure-function correlation we found in the Drosophila central brain is higher than typically reported in macro-scale networks in the human brain, where MRI-based structural and functional connectivity is similarly positively correlated, but generally with lower correlation coefficients (Honey et al. 2009) (for review, see Suárez et al. 2020). There are many factors that could contribute to this difference, including different levels of temporal and spatial precision in the functional measurements, vastly different amounts of biological detail in the structural connectivity measurements, and/or genuine biological differences between the two systems. Interestingly, the inclusion of detailed synaptic information (i.e. the number of synapses associated with a given inter-region connection) did not increase the correlation between structural and functional connectivity (Fig. 2). This suggests that synapse-level connectivity does not much constrain functional connectivity at the mesoscale level, despite synapse count being very important for smaller, functionally defined circuits (Takemura et al. 2017a; Takemura et al. 2017b). Mesoscale functional correlations reflect the influence of many neurons that individually belong to different functional circuits. While these individual neurons can vary substantially in their synapse number, when averaging over large brain regions, the synapse count connectivity is highly correlated with the cell count connectivity (r=0.90, data not shown).

In addition to the general correspondence between structural and functional connectivity, we found a number of other intriguing similarities between Drosophila central brain networks and networks in mammalian cerebral cortex. For example, we found that the structural network in the Drosophila central brain showed signs characteristic of small-world and scale-free networks (Fig. 3, and see Shih et al. 2015), which have been shown before in human structural and functional networks (Eguíluz et al. 2005; Sporns et al. 2004; Sporns and Zwi 2004). We also found that inter-region connectivity strengths are log-normally distributed (Fig. 1), which is also the case in mouse (Wang et al. 2012; Oh et al. 2014) and monkey (Markov et al. 2014) cortex. The observation that brain networks with such different biological and anatomical features share a key network topology suggests a deep correspondence in either developmental rules and/or functional constraints. It is not unusual for biological variables to be log-normally distributed. Indeed, multiplicative interactions among many variables will tend to produce log-normally distributed variables (Limpert et al. 2001; Buzsaki and Mizuseki 2014). In the context of nervous system development, interactions between pre- and post-synaptic cells that are shaped by cooperative combinations of adhesion and signaling molecules will be multiplicative. We speculate that the shared features between fly and vertebrate brains that can be observed at the mesoscale are the result of these universal phenomena at the microscale.

## Methods

### Data and code availability

All data and statistical analyses, modeling, and plot generation was performed using custom-written Python code, which can be accessed at https://github.com/mhturner/SC-FC. Data from the hemibrain connectome (v1.1) (Scheffer et al. 2020) was accessed using the Python neuPrint bindings. Functional data and derived structural connectivity measures can be accessed via figshare (https://doi.org/10.6084/m9.figshare.13349282).

### Analysis of connectome data to characterize inter-region structural connectivity

To compare brain regions in our atlas-aligned functional data to brain regions included and annotated in the hemibrain connectome, we used the mapping displayed in Supplemental Table 1. To measure structural connectivity from a source region, A to a target region, B, we used one of several connectivity metrics. (1) The primary metric used in the main text is the cell count connectivity metric. In this case, the connectivity weight from region A to B is defined as the total number of neurons that have at least one input synapse in region A and at least one output synapse (i.e. a presynaptic T-Bar) in region B. (2) A secondary metric we used was the T-Bar connectivity metric. In this case, we summed the number of T-Bars in region B that are associated with connecting cells from A (as defined in the cell count metric above). (3) The weighted synapse count is a metric defined in (Scheffer et al. 2020). This metric weights each connecting cell’s T-Bar count in region B by the fraction of total input synapses to that cell that are in region A. We measured the connectivity between each pair of regions included in this analysis (see Supplemental Table 1). To compare each region’s outgoing connectivity distribution to a log-normal distribution, we constructed a log-normal distribution for each region, based on that region’s mean and standard deviation connection weight. We then drew 1000 randomly-generated distributions (each with n=36, as in the data) from this fit model and compared this population of random distributions to the data using a Kolmogorov-Smirnov (KS) test. To combine distributions across regions, we first z-scored the log-transformed distribution for each region. This allows for different regions to have different means and standard deviations, but tests that they each follow their own log-normal distribution.

### Whole brain imaging to measure functional connectivity in the central brain

Details of whole brain imaging can be seen in (Mann et al. 2017). Imaging flies expressed GCaMP6s and myr::tdTomato pan-neuronally, and were of the following genotype:

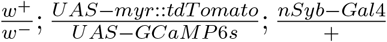

Flies were raised on molasses medium at 25°C with a 12/12 hr light/dark cycle. Flies were housed in mixed male/female vials and 5-day old females were selected for imaging. For imaging, flies were immobilized and their central brain exposed from the anterior of the head. Imaging experiments were performed on a resonant scanning two photon microscope (Bruker) equipped with a piezo Z drive which allows for fast volumetric acquisition. High resolution anatomical scans are taken as well as high speed functional scans at 3 *μm* isotropic spatial resolution and a temporal sampling frequency of 1.2 Hz. Anatomical scans are used to register to a common brain atlas, which is then used to extract time series data from each brain region.

### Statistical analyses and graph methods

We high-pass filtered each region-average fluorescence signal using a Butterworth filter with a cutoff frequency of 0.01 Hz to remove very slow drift in the fluorescence intensity. For display (Fig. 2) we converted each region response to a dF/F measurement relative to a baseline fluorescence defined as the mean fluorescence over the entire imaging session. To compute functional connectivity between a pair of regions, we used the Pearson linear correlation coefficient, r, between their two response traces. This is equivalent to the *δ*t=0 amplitude in the cross-correlograms shown in Fig. 2. To facilitate comparison across animals as well as to the structural connectivity, we Fisher z-transformed each inter-region correlation coefficient (see equation 1).

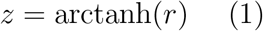

The functional connectivity matrix shown in Fig. 2 represents the average z-transformed correlation matrix across n=20 animals, but a connectivity matrix from an individual animal is also robustly correlated to the structural connectivity in the connectome (Fig. 2G).

To compare structural and functional connectivity (Fig. 2F), we first symmetrized the structural connectivity matrix by adding its own transpose to itself and dividing by 2. The functional connectivity matrix, by definition, is already symmetric. We then compared the upper triangle of the functional connectivity matrix to the upper triangle of the symmetrized structural connectivity matrix.

To compare graph features between the structural and functional networks, we used NetworkX (Hagberg et al. 2008) to create a graph for each network using its associated connectivity matrix. For the graph displays and analysis in Fig. 3, we first normalized each connectivity matrix by the maximum edge weight in the matrix. To compare the structural network to a matched random graph, we generated a binary adjacency matrix by applying a threshold to the structural connectivity matrix. We chose a threshold value of 50% of the maximum edge weight. For low threshold values, nearly every possible edge is realized and consequently little small world structure is observed. For high threshold values, the graph is no longer fully connected.

For the difference analysis in Fig. 4, we z-scored both the functional connectivity values as well as the (log-transformed) structural, cell-count connectivity values. This approach allowed us to better compare the structural and functional connectivity strength for a given connection, at least in relation to their respective connectivity distributions. We grouped brain regions by the Drosophila central brain super-region definitions in (Ito et al. 2014).

To perform the shortest path analysis in Fig. 4, we first defined the distance (D) for each connection as the inverse of that connection’s weight (W),(see equation 2).

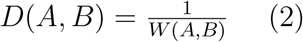

Using this distance definition, we used the Dijkstra’s Shortest Path First algorithm (Dijkstra 1959) in NetworkX to compute the shortest path between each pair of brain regions.

## Acknowledgments

We thank Estela Stephenson for technical support and members of the Clandinin lab for feedback on the work and on this manuscript. This work was supported by NIH grants F32-MH118707 (M.H.T.), NIH RO1EY022638 (T.R.C.), and a grant from the Simons Foundation (T.R.C.).

## Author contributions

M.H.T. and K.M. conceived the project. K.M. performed all the functional imaging experiments and registered *in vivo* data to the Drosophila brain atlas. M.H.T. analyzed functional connectivity data and structural connectivity data from the connectome, performed analyses and generated the figures. M.H.T., K.M. and T.R.C. wrote the manuscript. T.R.C. supervised all aspects of the work.

## Supporting Information

**Table 1:** Brain regions included in this analysis. Brain regions in the functional atlas, corresponding region(s) in the hemibrain connectome, super-region membership, and abbreviations. Super region abbreviations: AL-Antennal lobe; CX-Central complex; INP-Inferior neuropils; LH-Lateral horn; LX-Lateral complex; MB-Mushroom body; PENP-Periesophageal neuropils; SNP-Superior neuropils; VLNP-Ventrolateral neuropils; VMNP-Ventromedial neuropils.

| Atlas region | Connectome region(s) | Super-region | Abbreviation(s) |
| --- | --- | --- | --- |
| AL(R) | AL(R) | AL | AL: Antennal lobe |
| PB | PB | CX | PB: Protocerebral bridge |
| NO | NO | CX | NO: Noduli |
| EB | EB | CX | EB: Ellipsoid body |
| FB | FB, AB(R), AB(L) | CX | FB: Fan-shaped body; AB: Asymmetrical body |
| SCL(R) | SCL(R) | INP | SCL: Superior clamp |
| ATL(R+L) | ATL(R+L) | INP | ATL: Antler |
| ICL(R) | ICL(R) | INP | ICL: Inferior clamp |
| CRE(R+L) | CRE(R+L) | INP | CRE: Crepine |
| IB(R+L) | IB | INP | IB: Inferior bridge |
| LH(R) | LH(R) | LH | LH: Lateral horn |
| LAL(R) | LAL(R) | LX | LAL: Lateral accessory lobe |
| BU(R+L) | BU(R+L) | LX | BU: Bulb |
| MBCA(R) | CA(R) | MB | CA: Calyx |
| MBML(R+L) | $\beta L, \beta' L, \gamma L (R+L)$ | MB | ML: Medial lobe |
| MBPED(R) | PED(R) | MB | PED: Pedunculus |
| MBVL(R) | $\alpha L, \alpha' L (R)$ | MB | VL: Ventral lobe |
| CAN(R) | CAN(R) | PENP | CAN: Cantle |
| SMP(R+L) | SMP(R+L) | SNP | SMP: Superior medial protocerebrum |
| SLP(R) | SLP(R) | SNP | SLP: Superior lateral protocerebrum |
| SIP(R) | SIP(R) | SNP | SIP: Superior intermediate protocerebrum |
| AVLP(R) | AVLP(R) | VLNP | AVLP: Anterior ventrolateral protocerebrum |
| AOTU(R) | AOTU(R) | VLNP | AOTU: Anterior optic tubercle |
| WED(R) | WED(R) | VLNP | WED: Wedge |
| PLP(R) | PLP(R) | VLNP | PLP: Posteriorlateral protocerebrum |
| PVLP(R) | PVLP(R) | VLNP | PVLP: Posterior ventrolateral protocerebrum |
| GOR(R+L) | GOR(R+L) | VMNP | GOR: Gorget |
| VES(R) | VES(R) | VMNP | VES: Vest |
| SPS(R) | SPS(R) | VMNP | SPS: Superior posterior slope |
| EPA | EPA | VMNP | EPA: Epaulette |

**Figure S1:**
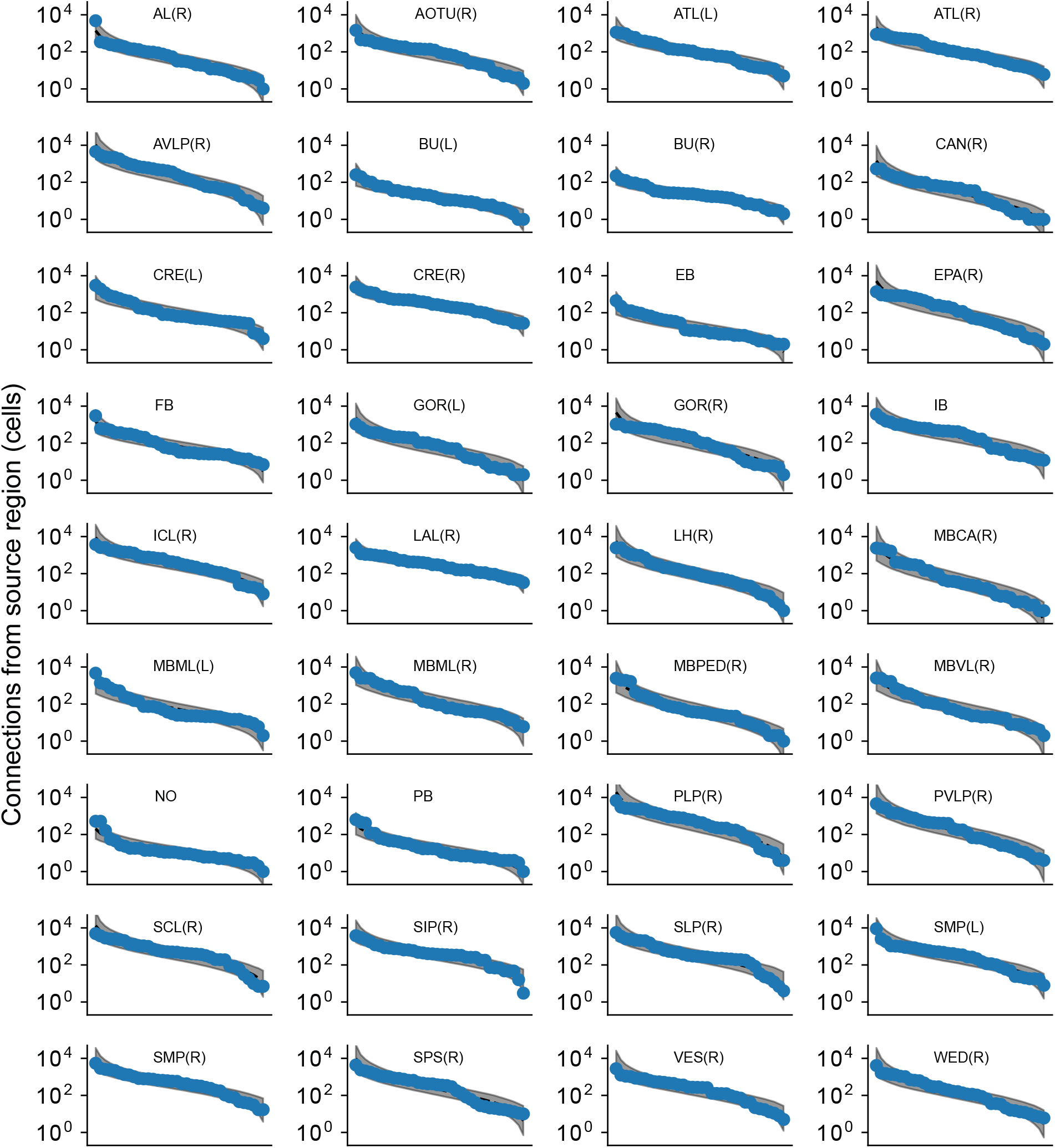
Distribution of outgoing connections from all brain regions. Each panel shows the ordered distribution of outgoing cell count connectivity from the indicated brain region (blue points). Gray shaded region indicates best-fit log-normal distribution.

**Figure S2:**
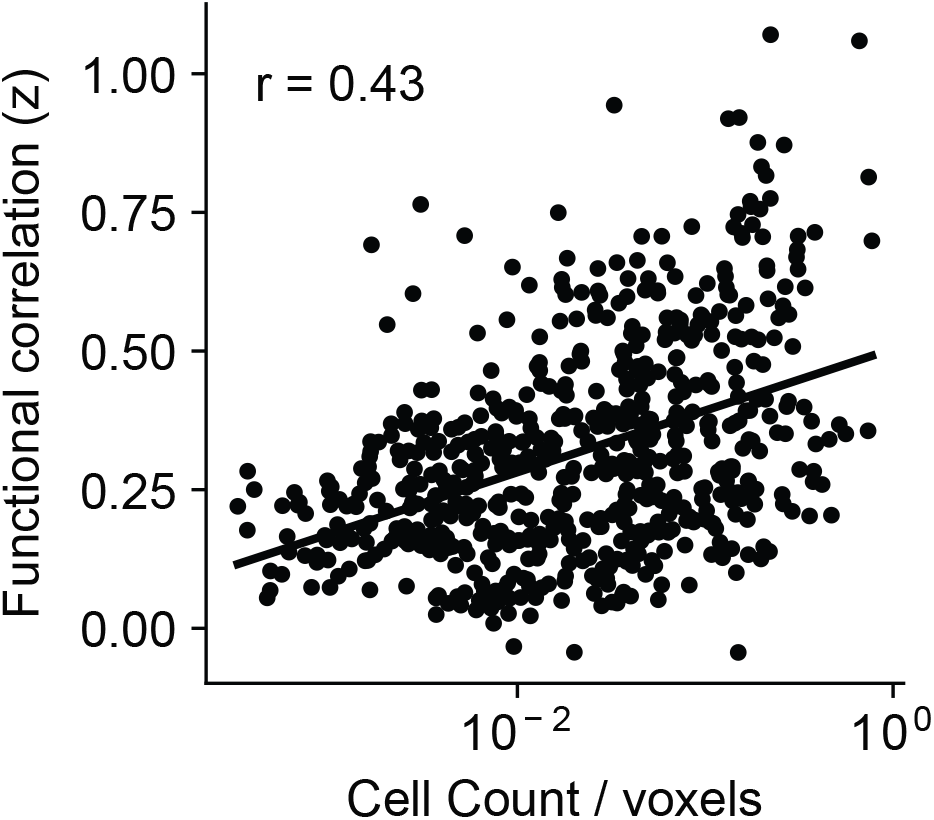
Volume-normalized cell count connectivity predicts functional connectivity. The cell count connectivity between each pair of regions (as in Fig. 2) was normalized by the geometric mean of the sizes (in voxels) of each region in the pair. This volume-normalized connectivity metric is still predictive of mesoscale functional connectivity.

**Figure S3:**
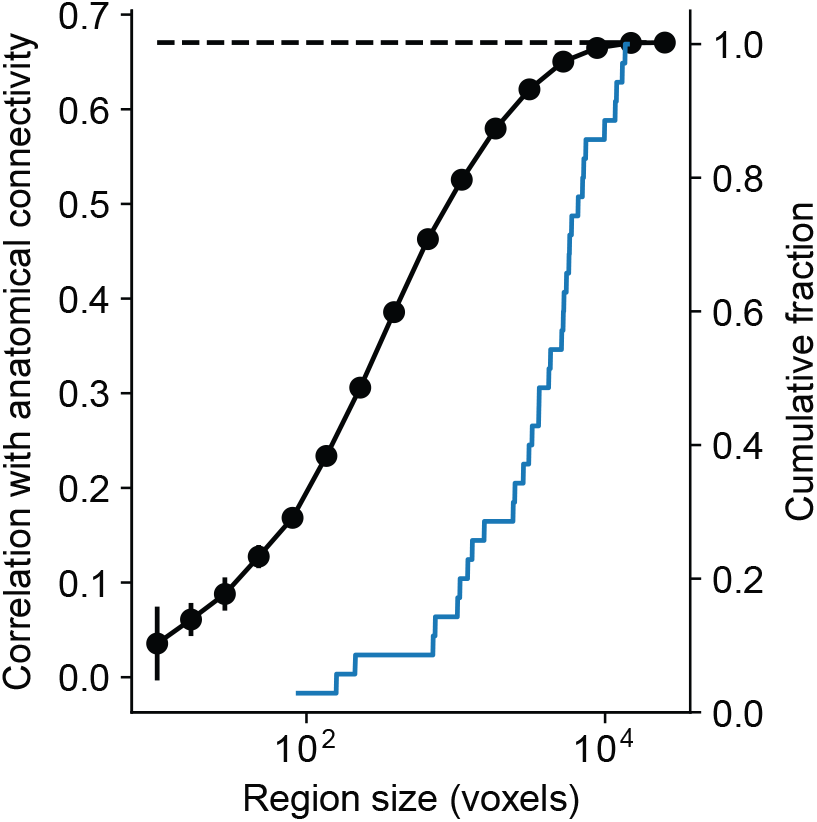
Region size does not preclude accurate measurement of structure function correlations. To assess whether relatively lower signal-to-noise in smaller brain regions limited our ability to infer functional connectivity and its correlation with structural connectivity, we performed a subsampling analysis. we randomly subsampled each region by the number of voxels indicated on the horizontal axis and measured structure-function correlations for each subsampled functional connectivity matrix. When only 100 voxels are included in each region, the estimated structure-function correlation is only 0.15, when up to 1000 voxels are included in each region the estimated structure-function correlation is 0.50. For comparison, the right axis shows the cumulative histogram of region sizes. The subsampled structure-function correlation is approximately 90% of the full estimate at the 30th percentile of region size, indicating that most of the regions are considerably larger than the sizes that interfere with the estimate of the functional connectivity.

**Figure S4:**
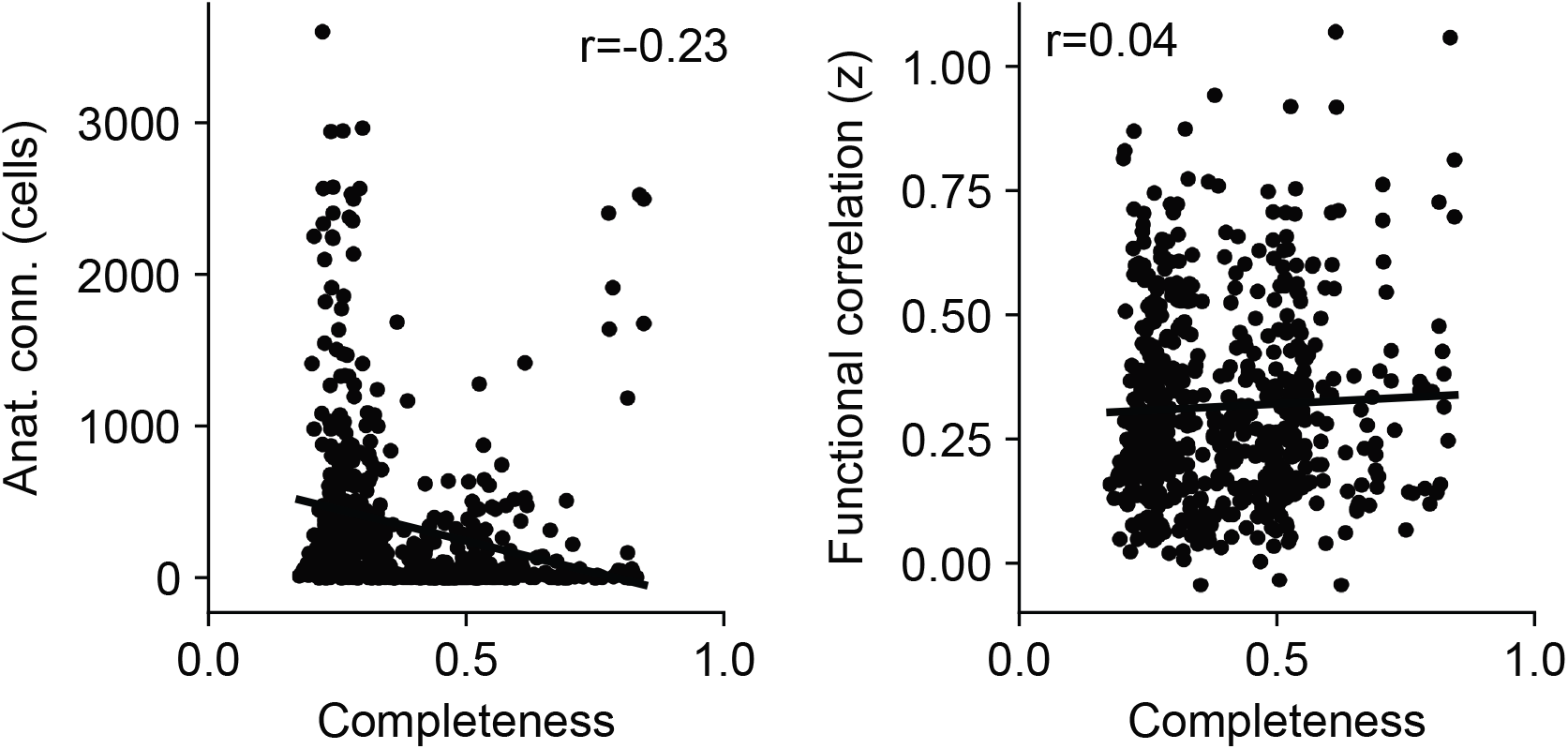
Relationship between reconstruction completeness connectivity. To understand the effect of the connectome reconstruction completeness on our estimates of structural connectivity, we computed a “completeness score” for each pair of brain regions, which was defined as the fraction of cell-assigned T-Bars in the source region multiplied by the fraction of cell-assigned postsynapses in the target region. Left: Higher completeness scores do not indicate stronger cell count connectivity, likely because this measure of connectivity is not sensitive to precise synapse numbers. Right: Higher completeness scores are not related to functional connectivity.

**Figure S5:**
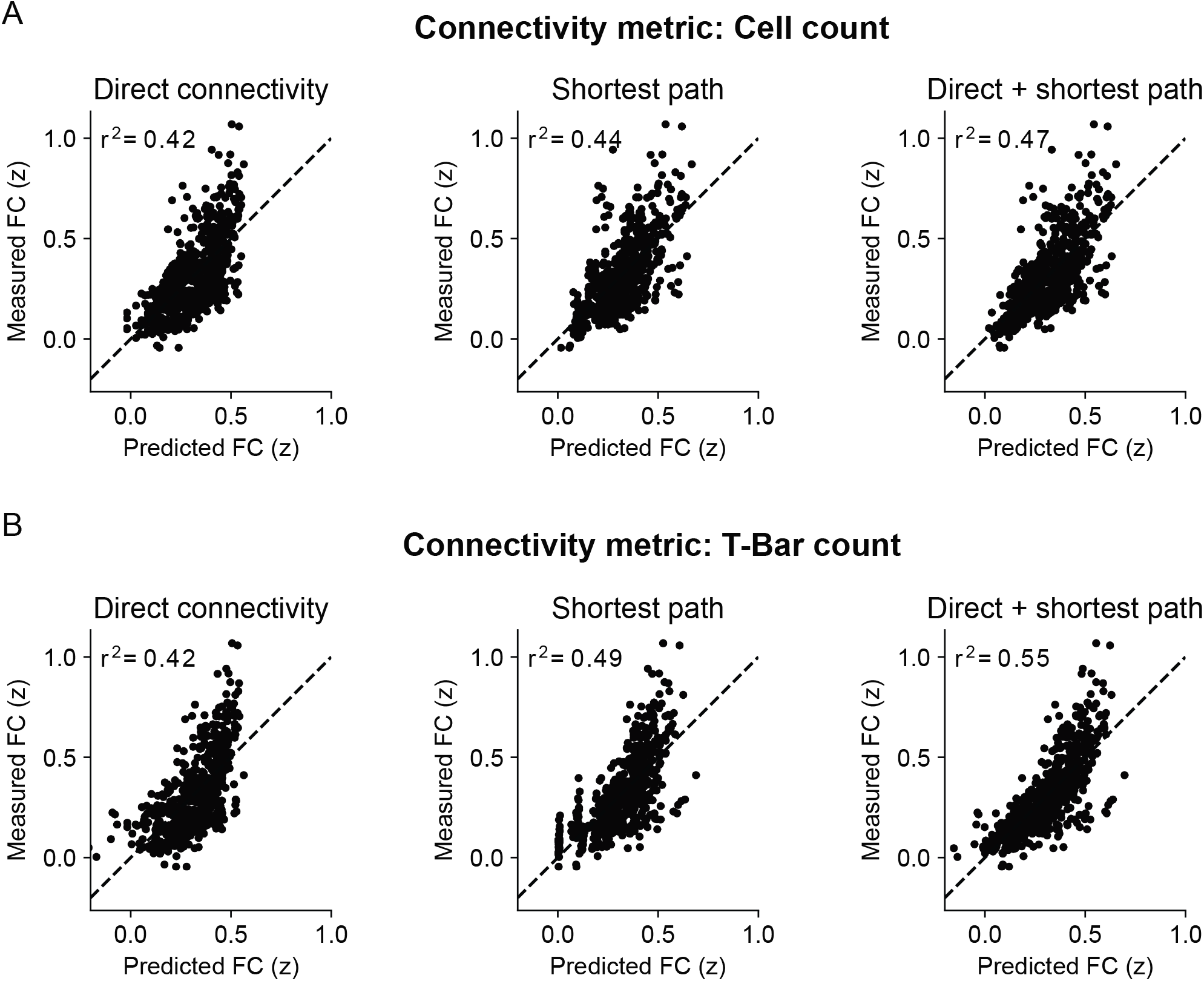
Linear regression models can account for roughly half of the variance in functional connectivity using structural connectivity. (A) Using the cell-count connectivity metric, we fit three linear regression models to predict measured functional connectivity based on: (i) Direct region-to-region connectivity (left); (ii) shortest path distance between each pair of regions (middle); and (iii) both direct connectivity and shortest path distance (right). For each model, the fraction of explained variance (*r*^2^), measured using 10-fold cross validation, is shown. (B) Same as (A) but using the T-Bar count connectivity metric instead of cell count connectivity.

## Notes

https://doi.org/10.6084/m9.figshare.13349282

https://github.com/mhturner/SC-FC

### Competing Interest Statement

The authors have declared no competing interest.

